# Comparative performance of the BGI and Illumina sequencing technology for single-cell RNA-sequencing

**DOI:** 10.1101/552588

**Authors:** Anne Senabouth, Stacey Andersen, Qianyu Shi, Lei Shi, Feng Jiang, Wenwei Zhang, Kristof Wing, Maciej Daniszewski, Samuel W Lukowski, Sandy SC Hung, Quan Nguyen, Lynn Fink, Ant Beckhouse, Alice Pébay, Alex W Hewitt, Joseph E Powell

**Affiliations:** Garvan-Weizmann Centre for Cellular Genomics, Garvan Institute of Medical Research, Darlinghurst, Sydney; Institute for Molecular Bioscience, University of Queensland, St Lucia, Brisbane; MGI, BGI-Shenzhen, Shenzhen 518083, China; BGI-Shenzhen, Shenzhen 518083, China; Menzies Institute for Medical Research, School of Medicine, University of Tasmania, Hobart; Department of Anatomy and Neuroscience, The University of Melbourne, Parkville, Melbourne; Department of Surgery, The University of Melbourne, Parkville, Melbourne; Centre for Eye Research Australia, Royal Victorian Eye and Ear Hospital, Melbourne; BGI Australia, 300 Herston Rd, Brisbane; Diamantina Institute, The University of Queensland, Woolloongabba, Brisbane; Centre for Eye Research Australia, The University of Melbourne, Royal Victorian Eye & Ear Hospital, Melbourne; St Vincent’s Clinical School, University of New South Wales, Sydney

## Abstract

The libraries generated by high-throughput single cell RNA-sequencing platforms such as the Chromium from 10x Genomics require considerable amounts of sequencing, typically due to the large number of cells. The ability to use this data to address biological questions is directly impacted by the quality of the sequence data. Here we have compared the performance of the Illumina NextSeq 500 and NovaSeq 6000 against the BGI MGISEQ-2000 platform using identical Single Cell 3’ libraries consisting of over 70,000 cells. Our results demonstrate a highly comparable performance between the NovaSeq 6000 and MGISEQ-2000 in sequencing quality, and cell, UMI, and gene detection. However, compared with the NextSeq 500, the MGISEQ-2000 platform performs consistently better, identifying more cells, genes, and UMIs at equalised read depth. We were able to call an additional 1,065,659 SNPs from sequence data generated by the BGI platform, enabling an additional 14% of cells to be assigned to the correct donor from a multiplexed library. However, both the NextSeq 500 and MGISEQ-2000 detected similar frequencies of gRNAs from a pooled CRISPR single cell screen. Our study provides a benchmark for high capacity sequencing platforms applied to high-throughput single cell RNA-seq libraries.

## Introduction

The human genome project was an important achievement in life sciences and paved the way for major technology developments in DNA and RNA sequencing. The development of synthesis-based Next-Generation Sequencing (NGS, also known as massively parallel or high-throughput sequencing) was pioneered by Solexa (1). After the company’s acquisition by Illumina, this technology was refined further and gave rise to a number of platforms that include the NextSeq, HiSeq and NovaSeq sequencers. These platforms have now produced the majority of the publicly available human sequencing data. Over time the cost of sequencing has decreased and the technology has become more accessible, both in terms of sequence hardware and tools for analysis (2). Collectively, this has resulted in NGS being adopted by many researchers, and used in clinical and industry settings.

Until recently, the majority of libraries sequenced have been generated on ‘bulk’ samples, consisting of the DNA or RNA collected from millions of cells. However, advances in single cell library preparation techniques (3, 4) have made it possible to produce sequencing libraries from tens of thousands of individually barcoded cells, and even individually barcoded molecules. High-throughput library preparation methods, such as the Chromium platform from 10x Genomics (5), are now widely available, enabling libraries consisting of tens of thousands of cells to be generated in several hours. The cDNA libraries from the Chromium experiments differ from ‘bulk’ libraries in that each cDNA molecule contains a Unique Molecular Identifier (UMI) and shared cell barcode. After amplification cDNAs are sheared, and adapter and sample indices are incorporated into finished libraries, which are compatible with next-generation short-read sequencing.

In 2017 BGI launched the MGISEQ-2000 as an alternative to existing short-read sequencing technologies (6). The technology underlying the MGISEQ-2000 combines DNA nanoball (DNB) nanoarrays (7) with polymerase-based stepwise sequencing (DNBseq), and its use has recently been validated as comparative in performance to the Illumina platforms when sequencing small noncoding RNAs (8), bulk transcriptomes (9), as well as whole genome DNA (10). To fully explore this platform’s potential for scRNA-seq, we undertook a direct performance comparison against Illumina technology by building scRNA-seq libraries generated with the Chromium platform from 10x genomics and sequencing 70,000 cells on both the MGISEQ-2000 and Illumina NextSeq 500, and NovaSeq 6000 (Figure 1).

**Figure 1.**
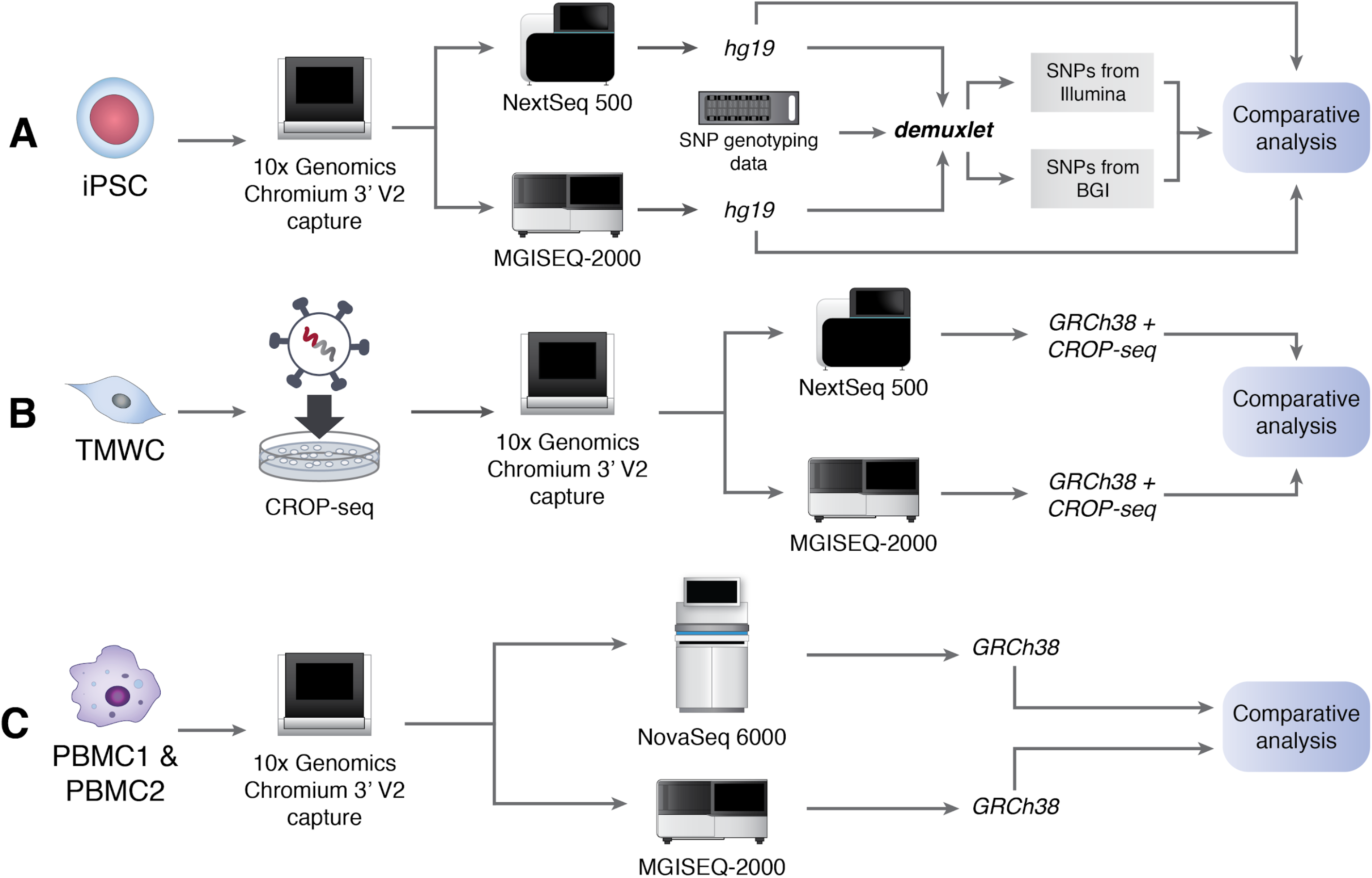
Experimental design. Preparation of single cell libraries and sequencing using Illumina and BGI platforms and subsequent analysis. A: Human **i**nduced **P**luripotent **S**tem **C**ells (**iPSC**) were generated generated from a human donor and underwent SNP genotyping in addition to scRNA-seq. B: **T**rabecular **M**esh**W**ork **C**ells (**TMWC**) derived from iPSCs were screened with a CRISPR-based molecular screen (CROP-seq). C: **P**eripheral **B**lood **M**ononuclear **C**ells (**PBMC**). Single cell libraries were prepared from two individual pools of PBMCs.

## Results

### Sequencing quality metrics

The total number of reads generated for the four libraries on the Illumina platforms was 159-616 million, and 1,112-1,339 million using the BGI platform. Comparison of sequencing quality control metrics revealed similar percentages of detectable valid cell barcodes, with a 0.8-1.2% greater detection from the NextSeq 500 and 1% from the NovaSeq 6000 (Table 1). The probability of a sequencing error is represented by a nucleotide base Q score, and thus the slightly higher percentages of valid cell barcodes from Illumina platforms most likely reflects the 5.1-6.8% (NextSeq 500) and 4.3-5.6% (NovaSeq 6000) higher Q30 scores observed in the cell barcode region of the reads compared to the MGISEQ-2000 (Table 1). A valid barcode is one that is detected from the sequence data that matches a whitelist of approximately 737,000 possible barcodes for the 3’ assay. The effect on the percentage of valid barcodes caused by lower Q30 score is partly mitigated by the Cell Ranger pipeline, which includes a step to correct for potential sequencing errors in the cell barcode based on a posterior probability that an observed barcode originated from the whitelist barcode. The second step in calling cells is based on the distribution and total counts of UMIs assigned to a given cell. We observed increased percentages of Q30 scores of 4.9-6.8% (NextSeq 500) and 4.1-5.9% (NovaSeq 6000) compared with the MGISEQ-2000 (Table 1).

**Table 1:** Sequence quality.

|  | iPSC |  |  | TMWC |  |  | PBMC1 |  |  | PBMC2 |  |  |
| --- | --- | --- | --- | --- | --- | --- | --- | --- | --- | --- | --- | --- |
|  | NextSeq<br>q 500 | MGISEQ-2000 | \|Δ\| | NextSeq<br>500 | MGISEQ<br>-2000 | \|Δ\| | NovaSeq<br>6000 | MGISEQ-2000 | \|Δ\| | NovaSeq<br>6000 | MGISEQ-2000 | \|Δ\| |
| Valid<br>barcodes | 97.20 | 96.40 | 0.80 | 97.90 | 96.80 | 1.10 | 98.00 | 97.00 | 1.00 | 98.00 | 97.00 | 1.00 |
| Reads<br>mapped<br>to<br>genome | 80.60 | 97.80 | 17.2 | 87.30 | 98.00 | 10.70 | 95.30 | 98.10 | 2.80 | 95.20 | 97.90 | 2.70 |
| Q30 in<br>barcode | 93.00 | 87.90 | 5.10 | 94.60 | 87.80 | 6.80 | 96.10 | 91.80 | 4.30 | 96.10 | 90.50 | 5.60 |
| Q30 in<br>UMI | 92.20 | 87.30 | 4.90 | 93.90 | 87.10 | 6.80 | 95.90 | 91.80 | 4.10 | 95.90 | 90.00 | 5.90 |
| Q30 in<br>RNA | 55.90 | 86.60 | 30.70 | 68.40 | 88.00 | 19.60 | 92.00 | 89.00 | 3.00 | 92.20 | 88.00 | 4.20 |
| Fraction<br>of reads<br>in cells | 79.20 | 80.10 | 0.90 | 95.00 | 95.10 | 0.10 | 93.70 | 94.80 | 1.10 | 94.10 | 95.20 | 1.10 |

The cell barcode and UMI enable individual cells to be identified for subsequent analysis, but these bases are obviously trimmed for the alignment stage. Accurate alignment to a reference transcriptome is partly a function of the sequencing error rate, and here we observe a dramatic difference in the Q30 percentage for the RNA transcript part of the read, with the MGISEQ-2000 achieving 19.9-30.7% greater Q30 compared to the NextSeq 500, although comparable performance compared with the NovaSeq 6000 (Table 1). The reduction in sequencing accuracy for scRNA-seq libraries on a NextSeq 500 has previously been discussed (11), and it has been hypothesised that this is due to flow cell surface chemistry. While variation in performance of specific flow cell lot numbers has been observed, it is important to note that the flowcells used here are not from a reported low-performance lot number.

The combination of assigning reads to a given cell, a transcript molecule, and aligning to a reference sequence directly affects the number of usable reads that are obtained from sequence data. Collectively, differences in the sequencing accuracy between platforms over the entire read length has consequential effects on the percentage of reads that pass quality control and that are able to be mapped to the reference genome. When we integrated the percentage of the reads that were able to be aligned to the GRCh38 (release 88) reference genome, we obtain an 11.1-17.2% difference between the NextSeq 500 and MGISEQ-2000, while the difference between the NovaSeq 6000 and MGISEQ-2000 is only 1.8-2.7% (Table 1). The lower percentage of alignment seen from the NextSeq 500 libraries is most likely due to the lower sequencing accuracy in the RNA transcript part of the read, as supported by the observation that the Q30 in RNA for Nextseq was much lower than that for MGISEQ-2000 (Table1). Because the thresholds used to determine if a read aligns to the genome are the same, the lower sequencing accuracy should not affect the biological interpretation of the aligned data. However, it does mean libraries sequenced on a NextSeq 500 will need to be sequenced at a greater depth to obtain the same sequencing depth of aligned reads per cell.

### Identification of cells, genes, and transcript molecules

To evaluate the similarity in the ability of sequencing platforms to identify the same cells, transcript molecules, and genes, we standardised the read depth between samples by downsampling. As the same cells from each sample had been sequenced on two platforms, we evaluated cell identification based on the observation of same cell barcode. Each of the two platforms identified close to 100% of cells in common in the four samples (Figure 2a). For cells identified by only one platform, the mean number of UMIs are on average one log2 lower than the cells identified as common between platforms (Figure 2b). There is a lower concordance of shared genes for these cells, suggesting that these ‘platform specific’ cells are possibly cell free transcripts that have not been adequately detected during quality control filtering by the cell singlet detection algorithm. An alternative explanation is that these are cells with low transcriptional abundance, although we observe no evidence for this scenario.

**Figure 2:**
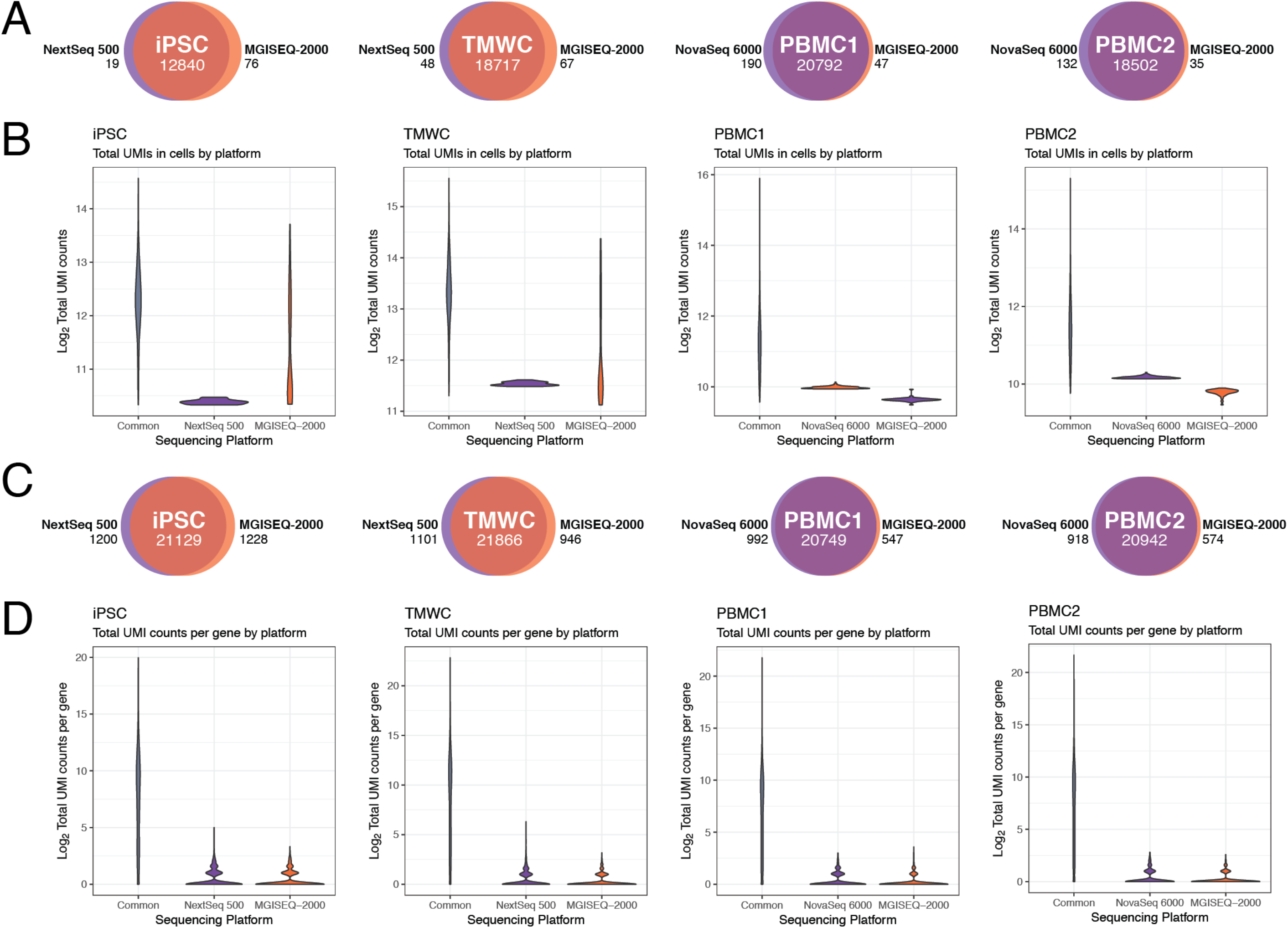
Cell barcodes and UMIs detected by platforms. We observed a high concordance of cell barcodes detected in common to both platforms (A), and the distributions of the Total UMI counts associated with a cell barcode shows a low average UMIs for cell barcodes detected by one platform (B). In each sample, both platforms detected similar total numbers of genes (C), although the mean of the UMIs for genes detected by a single platform shows that platform bias in gene detection was limited to lowly expressed genes (D).

Gene detection was similarly at high concordance with 89.7-93.3% of genes detected by both platforms for the four samples (Figure 2c). For all samples a subset of genes were detected by a single platform. The percentages of genes detected by a single platform were approximately equal for NextSeq 500 vs MGISEQ-2000, while the NovaSeq 6000 identified an additional 1.5-1.9% of genes. Details of the the genes detected from each platform are provided in Supporting Material Tables 1-4. Based on the number of UMIs, genes detected by a single platform were very lowly expressed, and variation in detection is expected due to the level of expression. To further confirm this, we downsampled to an average of 10^5^ reads per sample and repeated the comparison of gene detection. Interestingly, the NextSeq 500 detected an additional 0.8-1.7% genes in the iPSC and TMWC datasets, while the MGISEQ-2000 detected an additional 0.2-0.6% genes in the PBMC datasets.

The capture efficiency in gene detection levels, based on the relationship between the mean UMIs per gene and the number of genes detected was a similar for the iPSC and human trabecular meshwork cells (TMWC) samples sequenced on the NextSeq 500 and MGISEQ-2000. However, we observed a slight increase in the capture efficiency for the two PBMC samples sequenced on the NovaSeq 6000 in comparison to the MGISEQ-2000 (Figure 3a). This is likely a function of the slightly higher sequencing accuracy in the UMI region of the read (Table 1), corresponding to an increase in the mean UMIs per cell from the NovaSeq 6000 (Table 2). As expected we observe a marginal increase in the estimated dropout rate for the two PBMC samples from the MGISEQ-2000 compared with the NovaSeq 6000 (Figure 3b), although the correlation across all cells was high (0.989 and 0.988 respectively). Interestingly, there is no mean difference in the estimated dropout rate for the iPSC and TMWC samples between NextSeq 500 and MGISEQ-2000 (Figure 3b), and while the correlations across cells is lower (0.988 and 0.954 respectively) this is likely a function of the lower read depths for these samples, combined with greater variation in sequencing quality between platforms (Table 1). However, taken together, our analyses show that the gene detection, and quantification of transcript molecules via UMIs is highly consistent across platforms (Figure 3c-d).

**Figure 3.**
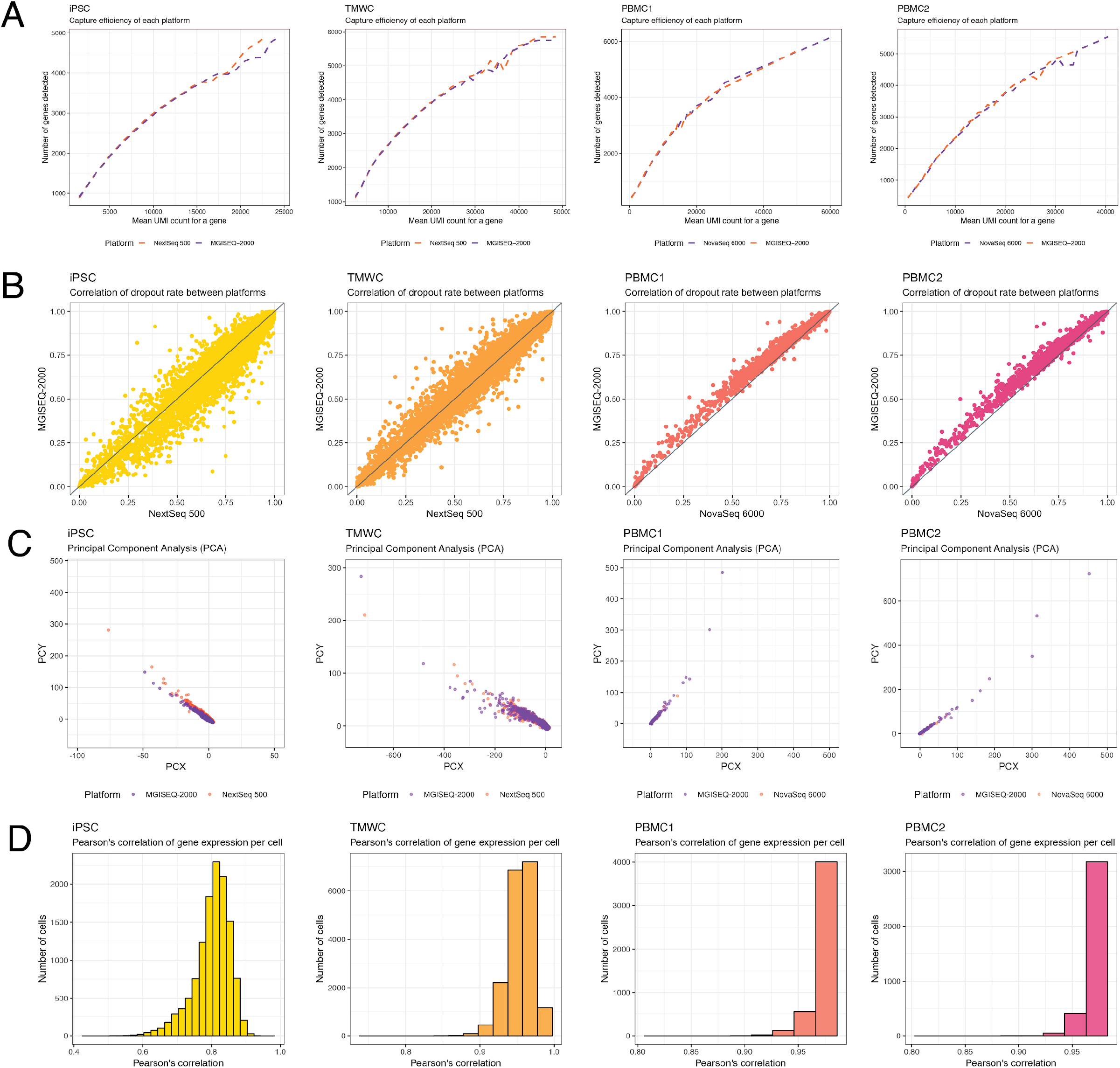
Comparison of gene capture between platforms. A: Capture efficiency of each platform. Efficiency is evaluated from the number of genes with the mean number of transcripts of a gene expressed by a cell. B: Correlation of dropout rate between platforms. Dropout rates for each platform were calculated by the DANB model from M3Drop package. C: Principal Components one and two calculated from 1,500 of the most variable genes. D: Correlation of gene expression in cells identified by both sequencing platforms. Correlation of each cell is represented in the histograms, while the expression values of genes in the cells with the lowest and highest correlations are represented in the scatter plots.

**Table 2:** Cell metrics.

|  | iPSC |  |  |  | TMWC |  |  |  |
| --- | --- | --- | --- | --- | --- | --- | --- | --- |
|  | NextSeq 500 | MGISEQ-2000 | MGISEQ-2000 (Equalised) | \|Δ\| (Equal depth) | NextSeq 500 | MGISEQ-2000 | MGISEQ-2000 (Equalised) | \|Δ\| (Equal depth) |
| Estimated number of cells | 12,859 | 12,916 | 12,916 | 57 | 18,765 | 18,784 | 18,784 | 19 |
| Total number of reads | 159,010,774 | 1,122,883,312 | 159,715,620 | 704,846 | 410,550,815 | 1,119,142,907 | 410,966,507 | 415,692 |
| Mean reads per cell | 12,365 | 86,937 | 12,366 | 1 | 21,878 | 59,579 | 21,879 | 1 |
| Median UMI counts per cell | 4,677 | 22,677 | 5,309 | 632 | 10,468 | 18,411 | 10,011 | 457 |
| Median genes per cell | 1,857 | 4,691 | 2,000 | 143 | 2,754 | 3,781 | 2,667 | 87 |
| Total number of genes detected | 22,329 | 25,154 | 22,357 | 28 | 22,967 | 23,999 | 22,812 | 155 |
|  | PBMC1 |  |  |  | PBMC2 |  |  |  |
|  | NextSeq 500 | MGISEQ-2000 | MGISEQ-2000 (Equalised) | Δ (Equal depth) | NextSeq 500 | MGISEQ-2000 | MGISEQ-2000 (Equalised) | Δ (Equal depth) |
| Estimated number of cells | 12,859 | 12,916 | 12,916 | 57 | 18,765 | 18,784 | 18,784 | 19 |
| **Total number of reads** | 159,010,774 | 1,122,883,312 | 159,715,620 | 704,846 | 410,550,815 | 1,119,142,907 | 410,966,507 | 415,692 |
| **Mean reads per cell** | 12,365 | 86,937 | 12,366 | 1 | 21,878 | 59,579 | 21,879 | 1 |
| **Median UMI counts per cell** | 4,677 | 22,677 | 5,309 | 632 | 10,468 | 18,411 | 10,011 | 457 |
| **Median genes per cell** | 1,857 | 4,691 | 2,000 | 143 | 2,754 | 3,781 | 2,667 | 87 |
| **Total number of genes detected** | 22,329 | 25,154 | 22,357 | 28 | 22,967 | 23,999 | 22,812 | 155 |

### Identification of genetic variation and CRISPR guides

The ability to call single nucleotide polymorphisms (SNPs) from scRNA-seq data allows researchers to use multiplexing strategies in the library generation stage, reducing the overall cost of running experiments where large sample sizes are needed (12). The power of demultiplexing a cell back to an individual donor is partly a function of the number of SNPs that can confidently be called from the short RNA section of the read. Using the iPSC sample, which comprises of cells multiplexed from two unrelated donors, we assigned cells to an origin donor by calling SNPs from the equalised total reads of sequence data generated by the NextSeq 500 and MGISEQ-2000 using the demuxlet algorithm (12). Donor identity was confirmed using genotyped SNPs from an Illumina Global Screening array that had been imputed to the Haplotype Reference Consortium panel (13). Despite equalised read depths across platforms, we identified an additional 1,065,659 SNPs from the MGISEQ-2000 data. The additional SNPs allowed demuxlet to assign an additional 1,694 cells to the correct donor (Table 3, Figure 4), with the greater SNP detection likely due to the higher sequencing quality of the MGISEQ-2000 reads (Table 1). To verify that this was not a function of differences in the base-pair length of the RNA section of the read, we trimmed the BGI data to a total RNA-read length of 98 bp and re-called SNPs, and could still correctly identify an additional 1,663 cells.

**Figure 4.**
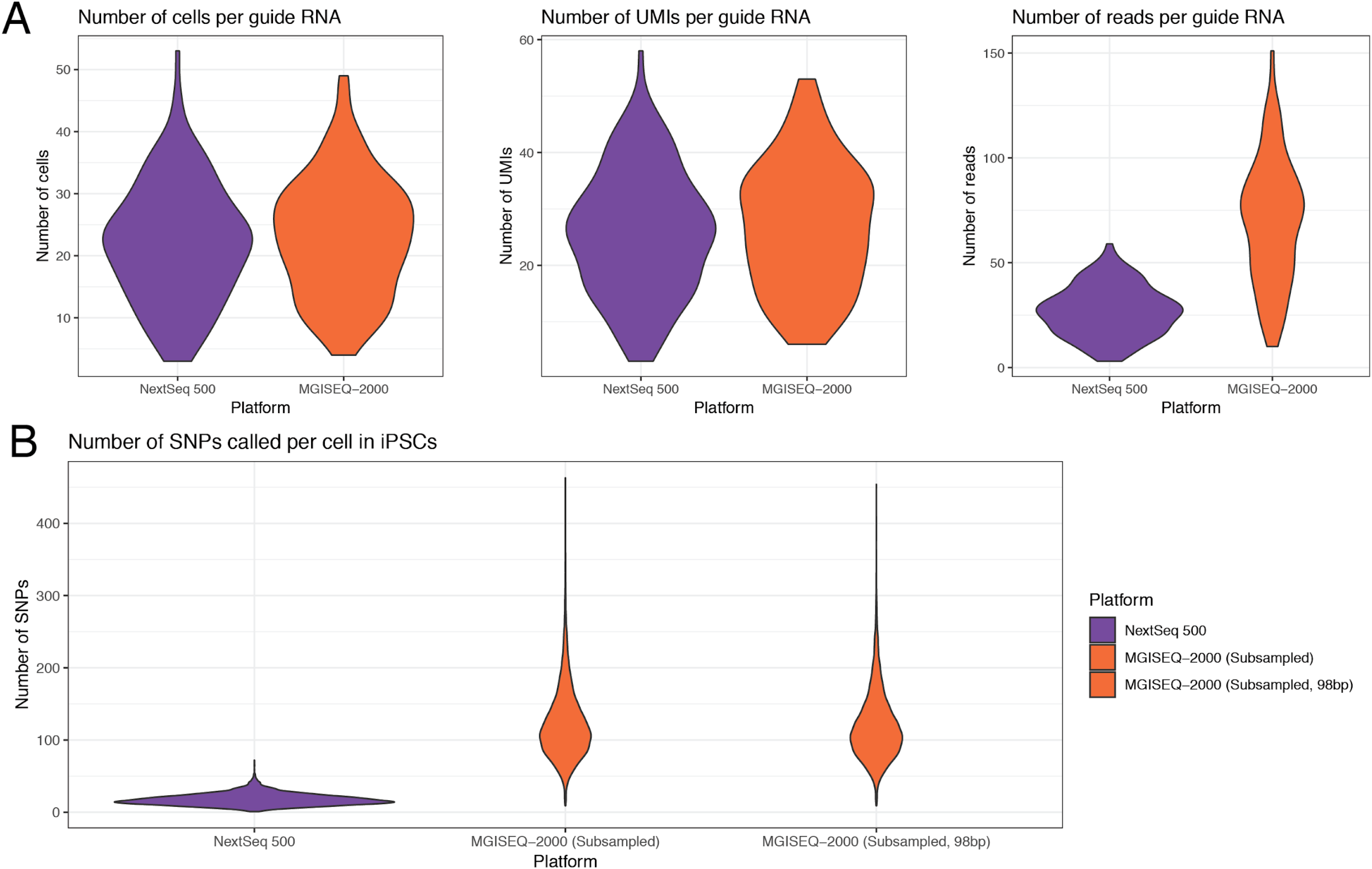
Experiment-specific metrics. A. Metrics related to guide RNA assignment in TMWC. This excludes cells that were not affiliated with a guide RNA and cells that with ambiguous assignments. B. Number of SNPs called per cell in iPSCs. SNPs were called from alignments of cells found in NextSeq 500 and MGISEQ-2000 datasets.

**Table 3:** Predicted assignments of cells to donor from iPSCs.

| **Prediction** | **NextSeq 500** | **MGISEQ-2000 (Downsampled)** | **MGISEQ-2000 (Downsampled,**<br>**98bp reads)** |
| --- | --- | --- | --- |
| **Unassigned** | 6016 | 4322 | 4311 |
| **Correctly assigned** | 4272 | 5966 | 5977 |

Finally, we evaluated the ability to detect the inserted guide RNAs (gRNA) from the TMWC that had been transfected with a CRISPR pool targeting 128 loci with the CROP-seq protocol. The guides are targeted to be inserted in the 3’ end of the gene and thus detectable from short-read sequence data. We observed consistent detection of the number of cells per guide, and the number of UMIs per guide across both the NextSeq 500 and MGISEQ-2000 (Figure 4).

## Discussion

To our knowledge, this study is the first to utilize MGISEQ-2000 platform for scRNA-seq, and the first to compare sequence performance for the widely used 10x Chromium platform against Illumina platforms. Our comprehensive benchmarking utilizes data from over 70,000 cells, and shows that the MGISEQ-2000 has to be highly comparable performance across a range of modalities to the Illumina NovaSeq 6000 platform, while being more cost effective (Supporting material table 5). When considering sequencing quality, as well as cell, UMI, and gene detection for single cell RNA-sequencing experiments, we found the Illumina NovaSeq 6000 and BGI MGISEQ-2000 platforms generated highly comparable data. Interestingly however, compared with the NextSeq 500, at equalised read depth, the BGI platform performs consistently better, identifying more cells, genes, and UMIs. We were able to call an additional 1,065,659 SNPs from sequence data generated by the BGI platform, enabling an additional one in seven cells to be assigned to the correct donor from a multiplexed library. It is also noteworthy that the NextSeq 500 and MGISEQ-2000 detected similar frequencing of gRNAs from a pooled CRISPR single cell screen. This work provides a benchmark for high capacity sequencing platforms applied to high-throughput single cell RNA-seq libraries.

## Methods

### Description of the single cell datasets and cell collection details

A total of four scRNA-seq libraries were generated from three experimental scenarios, chosen to evaluate the ability of sequencing platforms to provide sufficient information to detect features such as germline genetic variation and CRISPR inserts. All experimental work performed in this study was approved by the Human Research Ethics committee (HREC) of the Royal Victorian Eye and Ear Hospital (11/1031H; 13/1151H) or the Tasmanian Health and Medical HREC (H0012902) and conformed with the Declarations of Helsinki, under the requirements of the National Health & Medical Research Council of Australia (NHMRC).

#### iPSC

Consisted of undifferentiated human induced pluripotent stem cells (iPSCs) maintained with StemFlex (ThermoFisher Scientific) that were derived from two unrelated individuals (14). Colonies were harvested using ReleSR^TM^ (Stem Cell Tech) and were dissociated into a single cell suspension. Cells were counted and assessed for viability with Trypan Blue using a Countess II automated counter (Thermo Fisher Scientific), then pooled at a concentration of 391-663 cells/µL (3.91×10^5^ - 6.63×10^5^ cells/mL). Final cell viability estimates ranged between 95-97%. The two cell lines were then genotyped separately using the Infinium HumanCore-24 v1.1 BeadChip assay (Illumina), and SNPs were called from this assay with GenomeStudioTM V2.0 (Illumina). To generate the libraries, cells were partitioned and barcoded using high-throughput droplet 10x Genomics Chromium Controller (10x Genomics, USA) and the Single Cell 3’ Library and Gel Bead Kit (V2; 10x Genomics; PN-120237). The estimated number of cells in each well in the Chromium chip was optimized to capture approximately 10,000 cells. GEM generation and barcoding, cDNA amplification, and library construction were performed according to standard protocol.

#### TMWC

Comprised of cultured human trabecular meshwork cells (TMWCs) that had been transfected with a CROP-seq (Addgene: 99248) guide RNA (gRNA) pool targeting 128 loci, with the guides targeted to be inserted in the 3’ end of the the gene and thus detectable from short-read sequence data. TMWCs were plated in T75 flasks and transfected with a pooled single guide RNA (sgRNA) library lentivirus containing sgRNA for 128 targets, 10 of which were control genes. Cells were harvested 7 days after virus transduction and were FACS sorted for EGFP-positive and viable cells (propidium iodide-negative cells) before applying to the Chromium System (10x Genomics) single cell RNA-sequencing workflow. Single cell suspensions were used to generate a Chromium library using the Chromium Single Cell 3’ v2 Library (10x Genomics; PC-120237). The estimated number of cells in each well in the Chromium chip was optimized to capture approximately 10,000 cells.

#### PBMC1 and PBMC2

Consisted of peripheral blood mononuclear cell (PBMCs) collected from a total of 28 unrelated individuals. Peripheral blood samples were collected in Vacutainer Cell Preparation Tubes containing sodium heparin and ficoll (BD Biosciences: 362753), and were processed according to the manufacturer’s recommendations. Following separation, PBMCs were cryopreserved and stored. Samples were subsequently thawed, and each library contained a pool of PBMCs from 14 donors, with 40,000 cells loaded to achieve a targeted 20,000 cells per library.

### Illumina NextSeq 500 and NovaSeq 6000 sequencing

The iPSC and TMWC libraries were sequenced on an Illumina NextSeq 500 (NextSeq control software v2.0.2/ Real Time Analysis v2.4.11) using a 150 cycle NextSeq High Output Reagent Kit v2 in stand-alone mode as follows: 26bp (Read 1), 8bp (Index), and 98bp (Read 2). The NextSeq 500 sequencing was performed by the Institute of Molecular Bioscience sequencing core facility. The two PBMC libraries were sequenced on an Illumina NovaSeq 6000 (Software version: 1.4) using a 2×150 cycle S4 flowcell in standalone mode. The NovaSeq 6000 sequencing was performed by the Kinghorn Centre for Clinical Genomics Sequencing core facility.

### BGI MGISEQ-2000 sequencing

Libraries generated using the 10x Genomics Chromium system require a conversion step using the MGIEasy Universal Library Conversion kit (App-A) (Part Number: 1000004155) before sequencing can be performed on the MGISEQ-2000 instrument. For each library, 10ng was amplified using 10 cycles of PCR to incorporate a 5’ phosphorylation on the forward strand only. Purified PCR product was then denatured and mixed with a “splint” oligonucleotide that is homologous to the P5 and P7 adapter regions of the library to generate a circle (Figure S1). A ligase reaction was then performed to create a complete ssDNA circle of the forward strand then an exonuclease digest was performed to remove single stranded non-circularized DNA molecules. Circular ssDNA molecules then underwent Rolling Circle Amplification (RCA) to generate 300-500 faithful copies of the libraries which then fold upon themselves to become DNA Nanoballs (DNB). Each DNB library was then flowed across a 1,500M feature patterned array flow cell ready for sequencing using the MGISEQ-2000RS High-Throughput Sequencing Set (App-A) (PE100) (Part Number: 1000005662). The custom cycle mode on the instrument was run to allow 26bp (Read 1) and 100bp (Read 2) cycles without a index barcode read due to only one sample being run per flow cell, and FASTQ files were generated locally on the instrument. Sequencing was performed in BGI Shenzhen, MGI R&D facility.

### Bioinformatic and computational analysis

Sequencing data from both platforms were processed into transcript count tables using the Cell Ranger Single Cell Software Suite version 2.2.0 by 10x Genomics (http://www.10xgenomics.com/). Base calls from the NextSeq 500 and NovaSeq 6000 Illumina sequencers were pre-processed as described by Zheng et al. (5). Base calls from the MGISEQ-2000 were pre-processed as described by Huang et al. (15) into demultiplexed, processed reads. The BGI-formatted headers of the resulting FASTQ reads were converted to Illumina-formatted headers using custom Python scripts that are included with this publication’s accompanying repository. The quality of the raw sequencing data was assessed with FastQC v0.11.7 (16). The FASTQ files for both platforms were then processed with the *cellranger count* pipeline, where each sample was processed independently to generate the transcript count tables. Using STAR v2.5.1b (17), the iPSC library was mapped to the GRCh37/hg19 genome (release 84), while the PBMC libraries were mapped to the GRCh38 (release 88) *Homo sapiens* genome. The TMWC library was mapped to the GRCh38 (release 88) *Homo sapiens* genome that was spiked with gRNA and CROP-seq-associated sequences. This reference was prepared as described by Datlinger et al. (18). We note that, since the expression data is limited to the 3’ end of a gene and we used gene-level annotations, differences between reference versions, such as GRCh38, are unlikely to significantly alter conclusions. The resulting mapped counts for each pair of samples were then depth-equalized using the *cellranger aggr* pipeline, which downsampled raw reads from the higher-depth BGI library until the mean read depth per cell was equal to the mean read depth per cell of the Illumina library. Downsampling of mapped data to 10^5^ reads per sample was performed with DropletUtils (19).

Post-processing and biological analyses were performed on each sample using depth-equalized data. Statistical analyses were performed in R, using the ascend (*20*), scran (21), biomaRt (22) and M3Drop (23) packages. First, the count matrices were loaded into R and separated by platform. Cell barcodes were extracted from the matrices and those detected by both platforms were identified. The genes of these cells were then compared in terms of identity and distribution. Counts from each platform underwent quality control separately. A cell quality matrix based on the following data types: library size (total mapped reads), total number of genes detected, percent of reads mapped to mitochondrial genes, and percent of reads mapped to ribosomal genes. Cells that had any of the four parameter measurements higher than 3x median absolute deviation (MAD) of all cells were considered outliers and removed from subsequent analysis (Table S1). Next, we applied two thresholds to remove cells with mitochondrial reads above 20% or ribosomal reads above 50% (Table S1). To exclude genes that were potentially detected from random noise, we removed genes that were detected in fewer than 0.1% of all cells. The data from both platforms were combined back into one dataset. The NBDrop function from the M3Drop R package was used on filtered, un-normalised UMI counts to compare dropout rates between platforms. Abundantly expressed ribosomal protein genes and mitochondrial genes were then discarded to minimize the influence of those genes in driving clustering and differential expression analysis. Cell-cell normalization was performed using the deconvolution method described by (21). The correlation of gene expression between platforms was calculated using normalised UMI counts. To evaluate capture efficiency and transcript length bias of genes, gene lengths were calculated by summing exonic lengths retrieved from from the ENSEMBL *Homo sapiens* gene database. These values were then plotted as shown in Figure S2.

Additional analyses were conducted on the iPSC and TMWC samples to evaluate the influence of sequencing platform on properties specific to these experiments. Using genotype information from that was generated as described in (14), SNPs were called from the iPSC sample using *demuxlet* (12). To account for the downsampling of read depth in the MGISEQ-2000 data, only alignments from UMIs detected in the downsampled data were used. As the MGISEQ-2000 sequencer produced a longer insert read at 100bp, the iPSC sequencing data was re-mapped to the reference using reads that were truncated to 98bp. The reads were also downsampled to the same depth as the NextSeq 500 dataset. For the TMWC sample, gRNAs were detected using transcriptome data. This information was supplemented with read counts from the alignments using custom Python scripts that can be found in the accompanying repository.

## Data access

### Acknowledgments

We are grateful for the support in sample collection and processing performed by Antonia Rowson, Helena Liang, Linda Clarke and Qin Yi Lu. Illumina sequencing was performed by the Institute for Molecular Bioscience Sequencing Facility at the University of Queensland, and the Kinghorn Centre for Clinical Genomics Sequencing core facility. This work was supported by the Australian National Health and Medical Research Council (NHMRC) grants APP1132719, APP1083405 and APP1107599, Stem Cells Australia – the Australian Research Council Special Research Initiative in Stem Cell Science (J.E.P, A.W.H, A.P), The Macular Disease Foundation of Australia (A.P., A.W.H., J.E.P.), the Yulgilbar Alzheimer’s Research Program (J.E.P., A.P.). Further support was provided by a NHMRC Practitioner Fellowship 1103329 (A.W.H.), an NHMRC Senior Research Fellowship 1154389 (A.P.) and an Australian Research Council Future Fellowship FT140100047 (A.P.). Author contributions J.E.P. designed the study, acquired funding and led the analysis. A.S. performed bioinformatics, data processing, and computational analyses.

## Supporting material

### Supporting material tables

**Table S1:** Cell Filtering.

|  | Sample | Cells filtered by library size | Cells filtered by low expression | Cells filtered by mitochondrial transcript expression | Cells filtered by ribosomal transcript expression | Genes filtered by low expression |
| --- | --- | --- | --- | --- | --- | --- |
| Illumina NextSeq 500 | **iPSC** | 104 | 82 | 325 | 42 | 16578 |
|  | **TMWC** | 244 | 184 | 382 | 80 | 16264 |
| Illumina NovaSeq 6000 | **PBMC1** | 49 | 1 | 450 | 1 | 18483 |
|  | **PBMC2** | 64 | 0 | 382 | 0 | 18153 |
| MGISEQ-2000 | **iPSC** | 103 | 90 | 332 | 48 | 16413 |
|  | **TMWC** | 226 | 142 | 377 | 62 | 16439 |
|  | **PBMC1** | 57 | 2 | 432 | 1 | 18969 |
|  | **PBMC2** | 73 | 0 | 397 | 0 | 18668 |

### Supporting material figures

**Figure S1.**
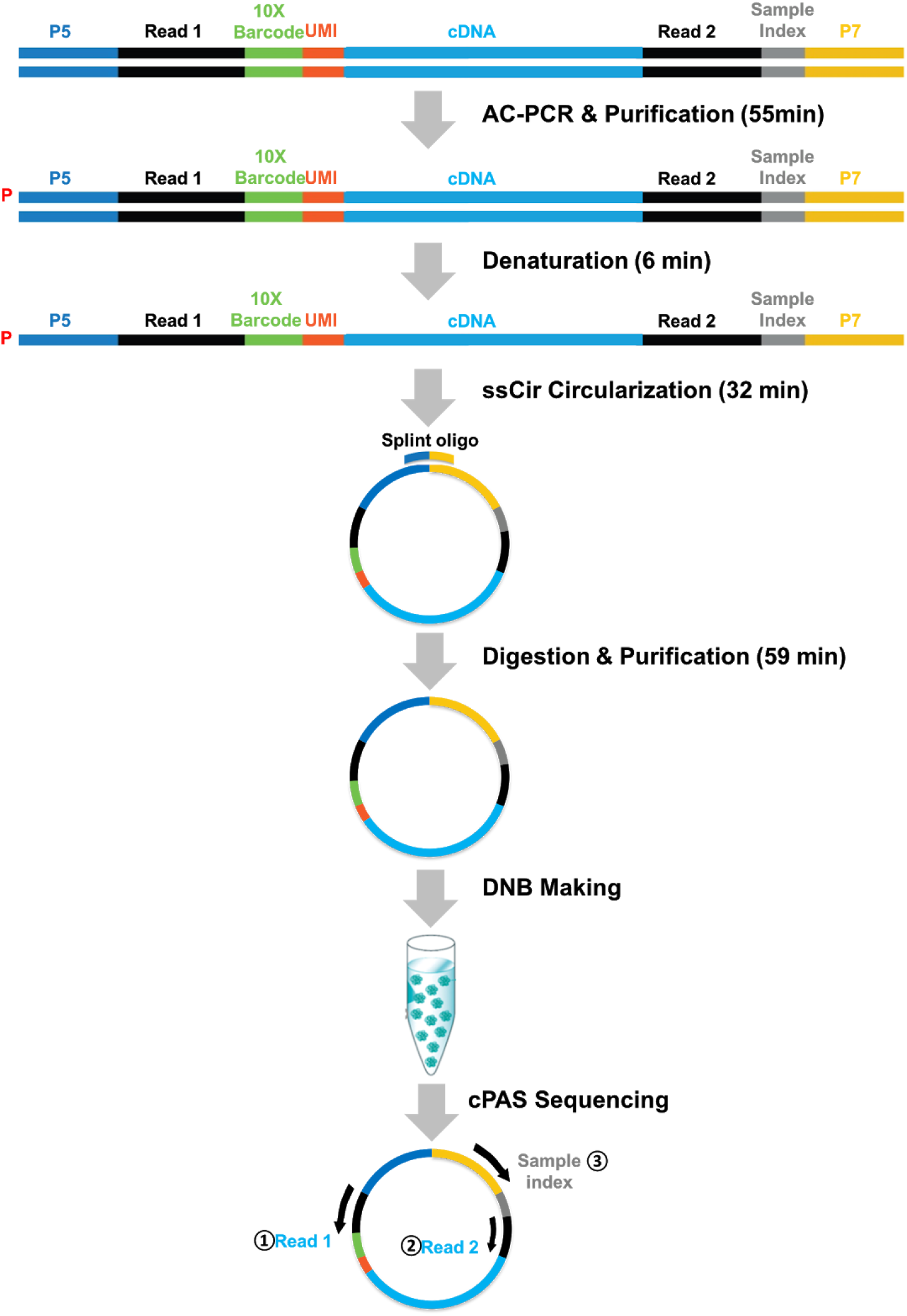
Conversion of Illumina-specific Single Cell library to BGI.

**Figure S2.**
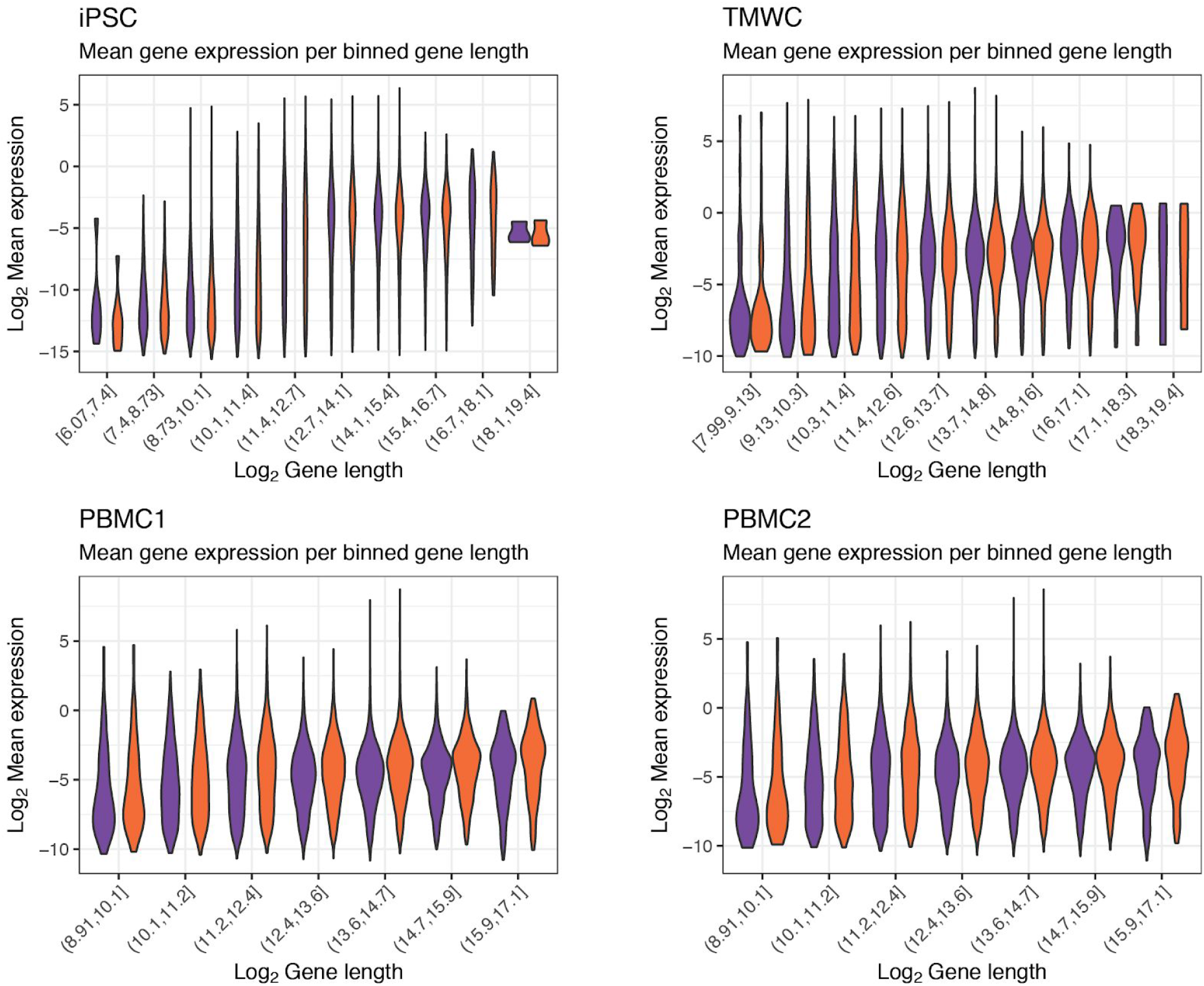
Relationship between gene length and abundance. Gene lengths were grouped into ten bins of equal number.

